# Palmitoylation of the oncogenic RhoGEF TGAT is dispensable for membrane localization and consequent activation of RhoA

**DOI:** 10.1101/062729

**Authors:** Jakobus van Unen, Dennis Botman, Taofei Yin, Yi I. Wu, Mark A. Hink, Theodorus W.J. Gadella, Marten Postma, Joachim Goedhart

## Abstract

Rho guanine exchange factors (RhoGEFs) control many aspects of the cellular cytoskeleton, and thereby regulate and control processes such as cell migration, cell adhesion and proliferation. TGAT is a splice variant of the RhoGEF Trio, with oncogenic potential. Whether the subcellular location of TGAT is critical for its activity is unknown. Confocal microscopy of fluorescent protein tagged TGAT revealed co-localization with a Golgi marker. Because plasma membrane localized RhoGEFs are particularly effective at activating RhoA, plasma membrane localization of TGAT was studied. In order to quantitatively measure plasma membrane association we developed a novel, highly sensitive image analysis method. The method requires a cytoplasmic marker and a plasma membrane marker, which are co-imaged with the tagged protein of interest. Linear unmixing is performed to determine the plasma membrane and cytoplasmic component in the fluorescence signal of protein of interest. The analysis revealed that wild-type TGAT is partially co-localized with the plasma membrane. Strikingly, cysteine TGAT-mutants lacking one or more palmitoylation sites in the C-tail, still showed membrane association. In contrast, a truncated variant, lacking the last 15 amino acids, TGAT^Δ15^, lost membrane association. The functional role of membrane localization was determined by measuring TGAT activity in single cells with a RhoA FRET-sensor and F-actin levels. Mutants of TGAT that still maintained membrane association showed similar activity as wild-type TGAT. In contrast, the activity was abrogated for the cytoplasmic TGAT^Δ15^ variant. Synthetic recruitment of TGAT^Δ15^ to membranes confirmed that TGAT effectively activates RhoA at the plasma membrane. Together, these results show that membrane association of TGAT is critical for its activity, but that palmitoylation is dispensable.

## Introduction

Rho GTPases are a subclass of the Ras superfamily of small GTPases, best known for their regulation of the cytoskeleton in eukaryotes (1, 2). Through remodeling of the F-­‐actin network they regulate several important cellular processes like cell migration, cell adhesion, proliferation and cell shape (3–5). Rho GTPases function as molecular switches that cycle between an active GTP-­‐bound form and an inactive GDP-­‐bound form (6). There are several classes of regulatory proteins that influence Rho GTPase activation cycle. Rho guanine exchange factors (RhoGEFs) activate Rho GTPases by accelerating the exchange of GDP for GTP (7). Rho GTPase activating or accelerating proteins (RhoGAPs) are responsible for turning Rho GTPases off by promoting the hydrolysis of the bound GTP to GDP (8). RhoGDIs sequester Rho GTPases in the cytoplasm in their inactive GDP bound state by binding to their prenylated C-­‐tails (9, 10). Deregulation of the RhoGTPase cycle has been mainly investigated within the context of cancer (11) and metastasis (12), but is also implicated in other pathologies like neurodegeneration (13), hypertension (14) and hemopathies (15).

The RhoGEF TGAT (*trio*-related transforming gene in ATL tumor cells) was first identified as an oncogenic gene product in adult T-cell leukemia cells (16). TGAT is formed by alternative splicing of the gene from the RhoGEF Trio, consisting of 255 amino acids encoding the second C-terminal RhoA activating Dbl homology (DH) domain of Trio and a unique extra 15 amino acid extension at its C-terminus. It was found that both its RhoGEF activity and the 15 amino acid extension were required for its transforming potential in NIH3T3 cells in vitro and in vivo (16), for the activation of tumorigenic transcription factor NF-κB via the IκB kinase complex (17), and the stimulation of matrix metalloproteinases (MMPs) via the inhibition of RECK (18). RhoGEFs have been put forward as possible therapeutic targets for the treatment of cancer (19, 20). TGAT is also considered as a possible therapeutic target for adult T-cell leukemia and several aptamer-derived inhibitors of TGAT were already developed (21). The 15 amino acid extension at the C-terminus of TGAT is referred to as the C-tail in this manuscript.

The molecular mechanisms underlying the contribution of the C-tail to the oncogenic potential of TGAT have not yet been investigated. Although Rho GTPases have been studied in detail for decades, the spatiotemporal aspects of their signaling and regulation are only starting to be uncovered (22). P63RhoGEF (23), a RhoA activating RhoGEF with a DH domain that has about 70% amino acid sequence homology to TGAT, is constitutively targeted towards the plasma membrane of cells by means of palmitoylation (24, 25). The activity of p63RhoGEF towards RhoA at the plasma membrane is auto-inhibited by a Pleckstrin Homology (PH) domain. A synthetic p63RhoGEF variant without the PH domain is constitutively active towards RhoA at the plasma membrane (26). Since TGAT consists of a DH domain, without an auto-inhibitory PH domain, it is likely that it has high basal RhoGEF activity. Because the C-tail is required for transforming activity, we hypothesized that the oncogenic potential of TGAT originates from subcellular targeting signals in the C-tail.

Here, we have used advanced fluorescence microscopy techniques in single living cells to investigate the role of several residues in the C-tail of TGAT on its subcellular location and function. A novel co-localization analysis based on confocal microscopy images shows that a TGAT mutant devoid of palmitoylation sites in its C-tail is still partially co-localized with the plasma membrane, whereas a TGAT mutant without the complete C-tail is exclusively located in the cytoplasm. Furthermore, we show that plasma membrane localization, but not palmitoylation of two cysteine residues in its C-tail, is necessary for actin polymerization and the activation of RhoA by TGAT. To confirm the plasma membrane as the subcellular site of action of TGAT GEF activity towards RhoA, we make use of a chemical heterodimerization system to target TGAT to several subcellular locations, and find that TGAT has the potential to activate RhoA on several endomembranes beside the plasma membrane.

## Results

### The influence of cysteines in the C-tail of TGAT on subcellular localization

The 15 C-terminal amino acid residues of TGAT have been described as essential for the oncogenic activity of TGAT (16). We set out to investigate the possible causes for this oncogenic activity in the C-tail in more detail. Close inspection of the sequence of the last 15 amino acid residues reveal two cysteines at position 242 and 253 that are putative palmitoylation sites (Figure 1A). This, in combination with the several basic and hydrophobic residues present in the C-tail, could potentially target TGAT to endomembrane structures, for instance the plasma membrane.

**Figure 1:**
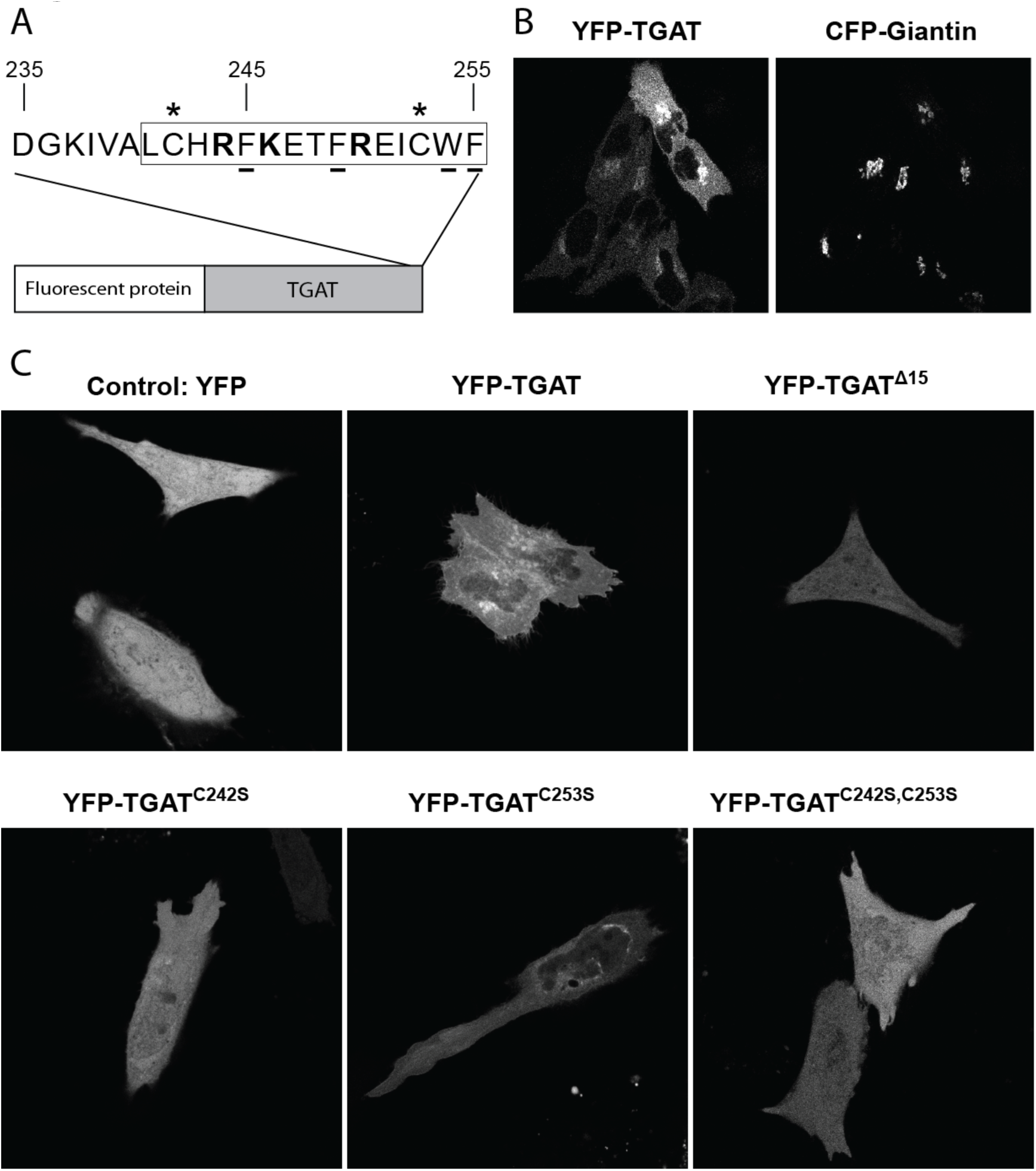
Subcellular localization of TGAT variants and quantification of their plasma membrane localization. (A) Schematic overview of TGAT fused to a fluorescent protein at its N-terminus. The blowout shows the last 20 amino acids of the C-tail of TGAT, with the last 15 amino acids indicated by the rectangle. Positively charged amino acids are shown in bold, amino acids with hydrophobic side-chains are underlined and the two cysteines at position 242 and 253 marked with an asterisk. (B) Confocal microscopy images of HeLa cells co-transfected with YFP-TGAT (*left*) and a golgi marker, CFP-Giantin (*right*). (C) Confocal microscopy images of HeLa cells transfected with YFP, YFP-TGAT, YFP-TGAT^Δ15^, YFP-TGAT^C242S^, YFP-TGAT^C253S^, YFP-TGAT^C242S, C253S^. Width of the individual images is 107µm in (B) and 110µm in (C).

In order to investigate the subcellular localization of TGAT, we constructed and visualized fusions of TGAT with a fluorescent protein attached to its N-terminus. HeLa cells transfected with YFP-TGAT and a golgi marker (CFP-Giantin) showed a strong co-localization of TGAT with the golgi apparatus, as previously observed for proteins that are palmitoylated (27, 28) (Figure 1B). To investigate the role of cysteine residues on the localization and function of TGAT, we constructed TGAT mutants by replacing the cysteine at amino acid position 242 (TGAT^C242S^) or 253 (TGAT^C253S^) or at both sites (TGAT^C242S, C253S^) by a serine. Confocal images were taken of HeLa cells transfected with a soluble YFP, YFP-TGAT, YFP-TGAT^Δ15^, YFP-TGAT^C242S^, YFP-TGAT^C253S^ or YFP-TGAT^C242S, C253S^ (Figure 1C). The golgi apparatus localization of TGAT was clearly diminished in the YFP-TGAT^Δ15^ and YFP-TGAT^C242S, C253S^ mutants, but still present to some extent in the YFP-TGAT^C242S^ and YFP-TGAT^C253S^ mutants. These results support the notion that palmitoylation of TGAT determines its subcellular localization.

### A new method to detect plasma membrane co-localization

It was previously shown that the isolated catalytic DH domain of p63RhoGEF at the plasma membrane is sufficient to induce constitutive GEF activity towards RhoA and induce actin remodeling (26). Since TGAT also consists of a constitutive active DH domain, its oncogenic potential might originate from localization at the plasma membrane. Because plasma membrane localization is not immediately apparent from the confocal images of TGAT (Figure 1B, C), we hypothesized that the fraction of TGAT at the plasma membrane might be very small compared to the unbound, intracellular component. In order to quantify the putative plasma membrane localization of TGAT and its mutants, we developed a novel co-localization method to analyze the confocal images of HeLa cells.

The method employs an untagged soluble RFP as marker for the cytoplasm and a lipid-modified CFP (Lck-CFP) as marker for the plasma membrane. The protein of interest with unknown localization is tagged with YFP, and its fluorescence is attributed to either the cytoplasm or the plasma membrane by linear unmixing. We briefly describe the procedure here, a detailed description of the quantification method can be found in the Material and Methods. Images of cells coproducing the three fluorescent proteins were used as input.

CFP, YFP and RFP channels were spatially registered and background subtracted (Figure 2A–C). A large number of perpendicular lines were carefully drawn on regions with a clean cytoplasmic-plasma membrane-extracellular transition in all possible orientations, from which average profiles were obtained using line-scans (normalized to cytoplasm) (Figure 2D-G). Subsequently, the average profiles were normalized to unity with respect to the cytoplasmic fluorescence level and, because the Lck-CFP marker is also localized in the cytoplasm, the profile was corrected by subtracting the normalized RFP profile (Figure 2G). The profiles from the YFP channel were subsequently unmixed into a cytoplasmic (CP) and plasma membrane (PM) component using constrained linear regression (Figure 2H). This procedure allowed for the detection of minute plasma membrane fractions.

**Figure 2:**
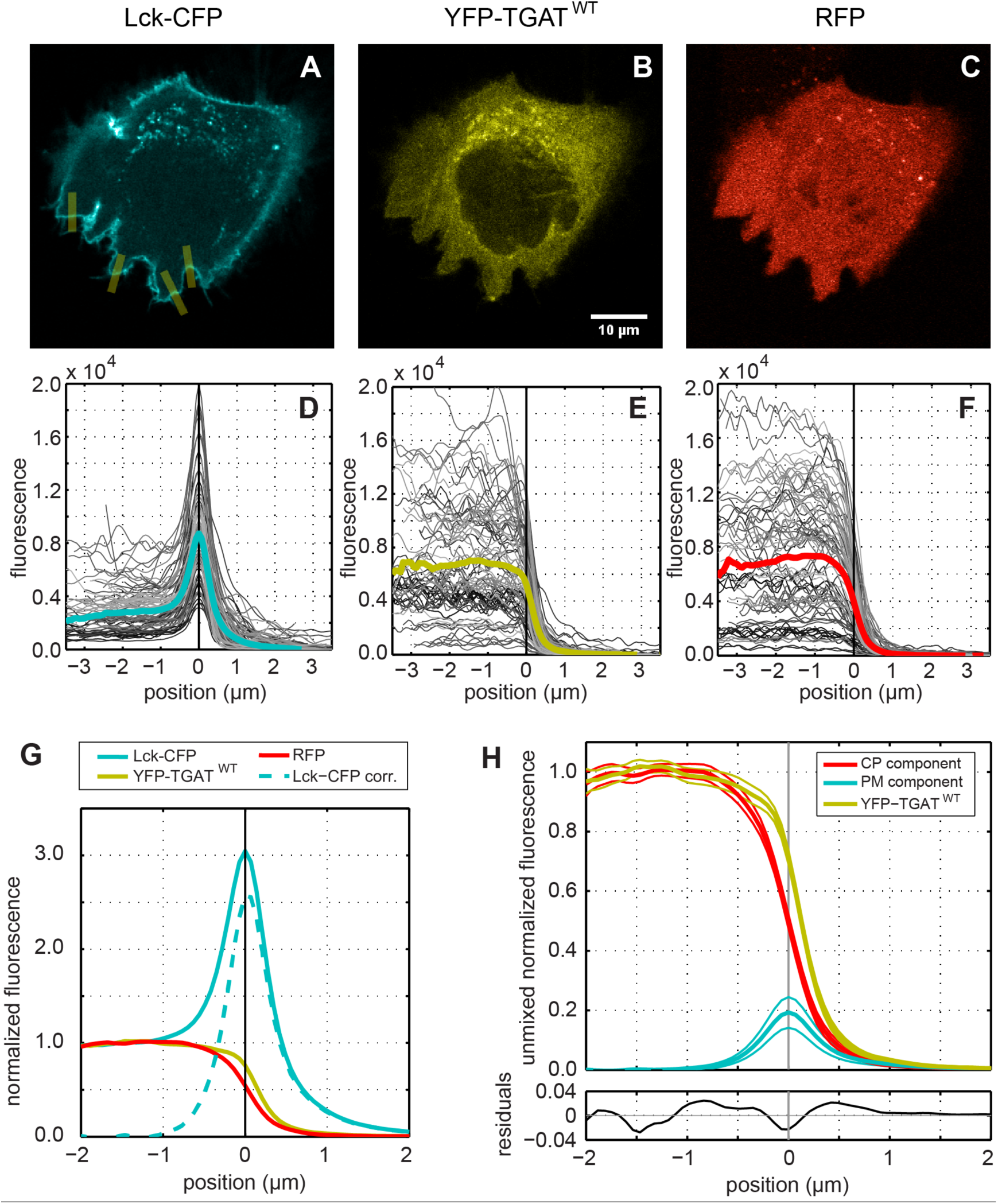
Quantification method for plasma membrane localization. High resolution confocal images (1024×1024 px with a pixel size of 140 nm) were taken of HeLa cells transfected with: (A) a plasma membrane marker Lck fused to CFP, (B) a protein of interest fused to YFP (wild type TGAT in this example) and (C) a cytoplasmic marker using a soluble RFP. The CFP channel was registered to the YFP channel based on the spatial shift determined with the positive control. The RFP channel was registered to the YFP channel based on the spatial shift determined with the negative control. Background fluorescence was subtracted from each image prior to processing. Using ImageJ many lines (10 px wide and 6-10 µm long) were carefully drawn perpendicular to the plasma membrane in regions with a well-defined cytoplasm-plasma membrane-extracellular space transition (yellow lines in panel A). For each channel line-scans were performed with the same lines using linear interpolation, the profiles were aligned and centered based on the peak in the Lck-CFP channel and placed on the same axis (panel D). The other channels were aligned and centered using the same positional shift and axis (panels E and F). Subsequently the average profiles were calculated (colored lines in panels D-G), and normalized to unity with respect to the cytoplasm fluorescence level (panel G). Because the Lck-CFP marker is also localized in the cytoplasm, the profile was corrected by subtracting the normalized RFP profile (dashed line panel G). In order to extract the cytoplasmic (CP) and plasma membrane (PM) component, the YFP profile was unmixed using the normalized RFP profile and the corrected Lck-CFP profile (panel H). The 95% confidence intervals (thin solid lines above and below the profiles) were estimated using bootstrapping (See material and Methods for details).

### C-tail palmitoylation is dispensable for plasma membrane localization of TGAT

To examine the plasma membrane localization of TGAT and its variants, we employed the novel co-localization analysis. First, we examined the dynamic range of our method by analyzing maximal and minimal plasma membrane association by employing a plasma membrane associated YFP (Lck-YFP) and a soluble YFP respectively. For the membrane bound positive control (Lck-YFP) we determined a PM-localized peak of 316 ± 20% (Figure 3A), whereas for the cytosolic negative control the PM localized peak was 100-fold lower, 3 ± 1% (Figure 3B).

**Figure 3:**
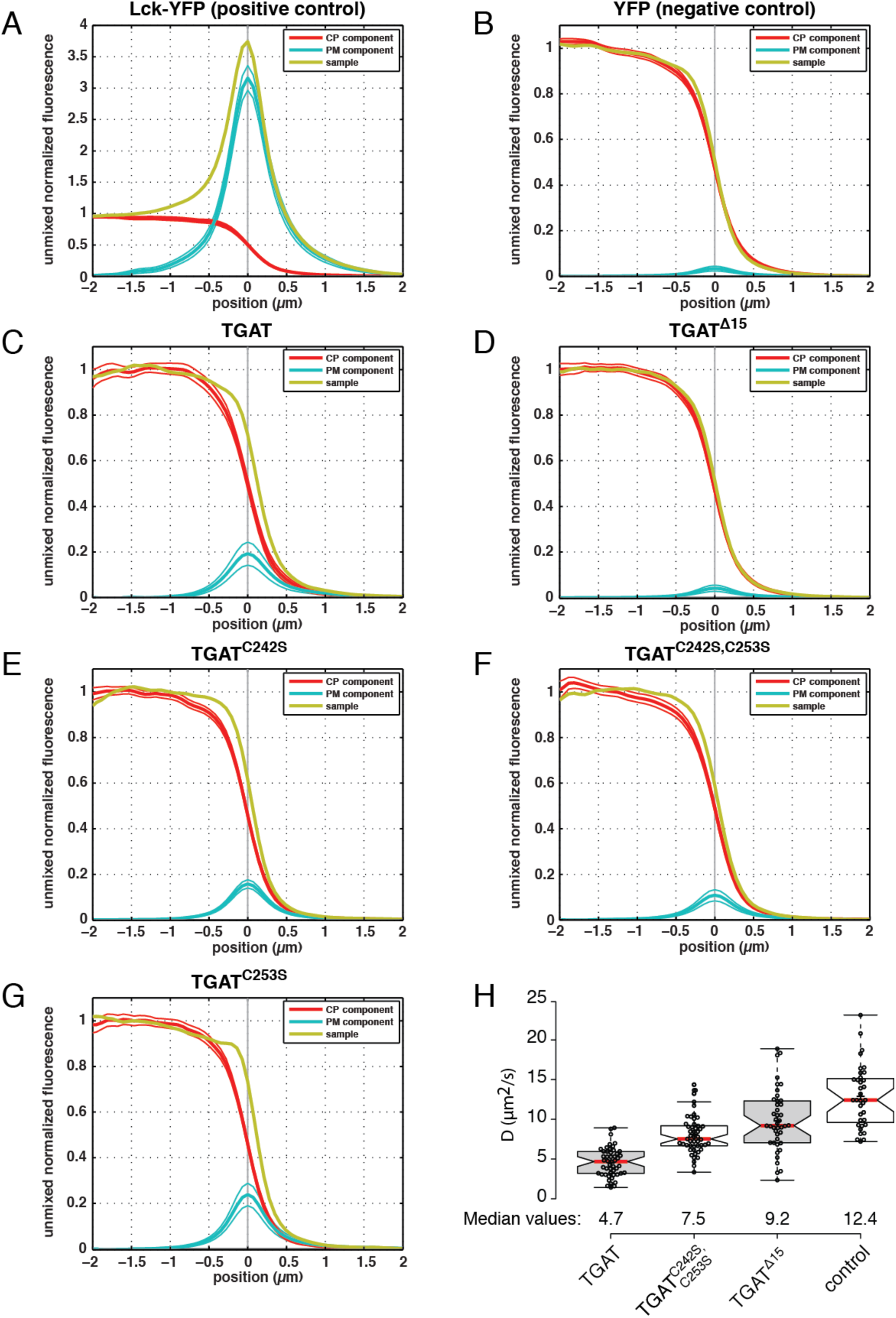
Determination of plasma membrane localization of all TGAT variants. Confocal microscopy was performed on HeLa cells transfected with a plasma membrane marker (Lck-CFP), a cytoplasm marker (soluble RFP) and either (A) Lck-YFP (*n* = 67 lines, 10 cells), (B) YFP (*n* = 66 lines, 12 cells), (C) YFP-TGAT (*n* = 88 lines, 30 cells), (D) YFP-TGAT^Δ15^ (*n* = 130 lines, 19 cells), (E) YFP-TGAT^C242S^ (*n* = 129 lines, 12 cells), (F) YFP-TGAT^C242S, C253S^ (*n* = 112 lines, 12 cells) or (G) YFP-TGAT^C253S^ (*n* = 79 lines, 5 cells). Averaged line profiles were obtained using a Matlab script and ImageJ, the profiles were normalized to cytoplasmic levels. Line profiles for the YFP channel (*yellow*) were subsequently unmixed in order to obtain the plasma membrane (*blue*) and cytoplasmic (*red*) component in each construct of interest. The solid thin lines above and below the PM and CP profiles represent the 95% confidence interval obtained from statistical bootstrapping. See Material and Methods for more details. (H)Diffusion coefficients of different TGAT mutants, as determined by fluctuation correlation spectroscopy (FCS). Diffusion coefficients were determined for HeLa cells transfected with CFP-TGAT (*n* = 49), CFP-TGAT^C242S, C253S^ (*n* = 49), CFP-TGAT^Δ15^ (*n* = 42) or free CFP (control, *n*= 35). Boxplot center lines represent the median values (*red*); box limits indicate the 25th and 75th percentiles as determined by R software; whiskers extend 1.5 times the interquartile range from the 25th and 75th percentiles; data points from individual cells are plotted as dots.

Next, we examined the plasma membrane association of TGAT and its variants, i.e. YFP-TGAT, YFP-TGAT^Δ15^, YFP-TGAT^C242S^, YFP-TGAT^C253S^ or YFP-TGAT^C242S, C253S^. Wild type TGAT (Figure 3C) clearly has a detectable PM component of about 19 ± 5%, which is significantly higher than the negative control. This indicates that TGAT is partly localized at the PM, albeit with a more then 10-fold lower level compared to the positive control Lck-YFP. The mutant TGAT^Δ15^, has a PM peak value of 4 ± 1%, which is statistically indistinguishable from the negative control, implying a full cytoplasmic localization (Figure 3D).

The mutants with the single point mutations, TGAT^C242S^ and TGAT^C253S^ (Figure 3E,G) exhibit PM peaks of 16 ± 2% and 24 ± 5% respectively. These levels are comparable to TGAT and suggest that these mutants are still localized at the plasma membrane at similar levels as TGAT. The variant with the double point mutation, TGAT^C242S, C253S^ exhibits a PM peak of 11 ± 3%, which is twofold lower than the TGAT level, but higher than the negative control at a 95% confidence level. This suggests that the double cysteine mutant of TGAT still associates with the plasma membrane.

Thus far, the results show that the cysteine mutants still have affinity for the plasma membrane. To independently verify this observation, we decided to measure the diffusional mobility of TGAT and its mutants. If the double mutant binds membranes, TGAT^C242S, C253S^ should diffuse slower through the cell then TGAT^Δ15^. To investigate this hypothesis, we employed fluorescence correlation spectroscopy (FCS) to measure and compare the mobility of different TGAT variants. HeLa cells were transfected with CFP-TGAT, CFP-TGAT^C242S, C253S^, CFP-TGAT^Δ15^ or just a soluble CFP and point scanning FCS measurements were performed in cell peripheries. Diffusion times were obtained by fitting the autocorrelation curves (see material and methods for details). From this, diffusion coefficients were calculated by correcting the measured diffusion time with the calibrated detection volume. Both CFP-TGAT (*D* = 4.7µm^2^/s, 95% CI [4.1 – 5.3]) and CFP-TGAT^C242S, C253S^ (*D* = 7.5 µm^2^/s, 95% CI [7.0 – 8.1]) exhibit a lower mobility than CFP-TGAT^Δ15^ (*D* = 9.2µm^2^/s, 95% CI [8.0 – 10.5]), although CFP-TGAT is still considerably less mobile than the double cysteine mutant (Figure 3H). Soluble CFP exhibits the highest mobility (*D* = 12.4µm^2^/s, 95% CI [11.0 – 13.9]) through the cell periphery.

From the mobility data we conclude that palmitoylation indeed has an effect on the plasma membrane residence time of TGAT, but the difference in diffusion time of TGAT^C242S, C253S^ when compared to that of TGAT^Δ15^ suggests that a fraction of TGAT^C242S, C253S^ is still interacting directly or indirectly with the plasma membrane.

### The influence of cysteines in the C-tail of TGAT on activation of the small GTPase RhoA

We next set out to explore the influence of plasma membrane affinity and palmitoylation of the C-tail of TGAT on its functional properties. In order to investigate the possible activation of RhoA by TGAT and its mutants, we transfected HeLa cells with the DORA RhoA biosensor and one of the TGAT variants. The YFP/CFP FRET ratio was used to assess the RhoA activation state in each condition (Figure 4). The minimal and maximal FRET ratios were estimated from the inactive non-binding RhoA_sensor_-nb bio_sensor_ (nb, 0.71, 95% CI [0.65 – 0.77]) and the constitutive active RhoA_sensor_-ca bio_sensor_ (ca, 6.12, 95% CI [5.62 – 6.62]), respectively. The values of the wild-type RhoA_sensor_-wt (wt) bio_sensor_ are expected to fall within this range of YFP/CFP ratios. Control cells transfected with the RhoA_sensor_-wt and RFP-RhoGDI show a decreased basal median ratio (0.56) compared to cells transfected with RhoA_sensor_-wt and a soluble RFP (0.86), illustrating the preserved regulation of the DORA RhoA bio_sensor_ by RhoGDIs. Cells transfected with the RhoA_sensor_-wt and RFP-TGAT^Δ15^ showed much lower basal median YFP/CFP FRET ratio (1.27, 95% CI [1.13 – 1.42]) when compared to the RFP-TGAT condition (3.27, 95% CI [3.13 – 3.41]). Cells transfected with the RhoA_sensor_-wt and RFP-TGAT^C242S^ (3.25, 95% CI [3.11 – 3.39]), RFP-TGAT^C253S^ (3.24, 95% CI [3.07 – 3.41]) or RFP-TGAT^C242S, C253S^ (3.31, 95% CI [3.18 – 3.44]) did not show a difference in basal median YFP/CFP FRET ratio compared to the RFP-TGAT condition. In order to directly test the influence of palmitoylation on the basal YFP/CFP FRET ratio, cells transfected with the RhoA_sensor_-wt and wild-type TGAT were incubated overnight in medium containing 25µM 2-bromo-palmitate (2-BP). No difference in basal median YFP/CFP FRET ratio was observed for 2-BP incubated cells (3.36, 95% CI [3.24 – 3.48]) when compared to the untreated cells (3.27, 95% CI [3.13 – 3.41]). Taken together, these results show that the plasma membrane localization conferred by the C-tail contributes to RhoA activation, but suggest that palmitoylation of the C-tail of TGAT does not influence the activation of RhoA in HeLa cells.

**Figure 4:**
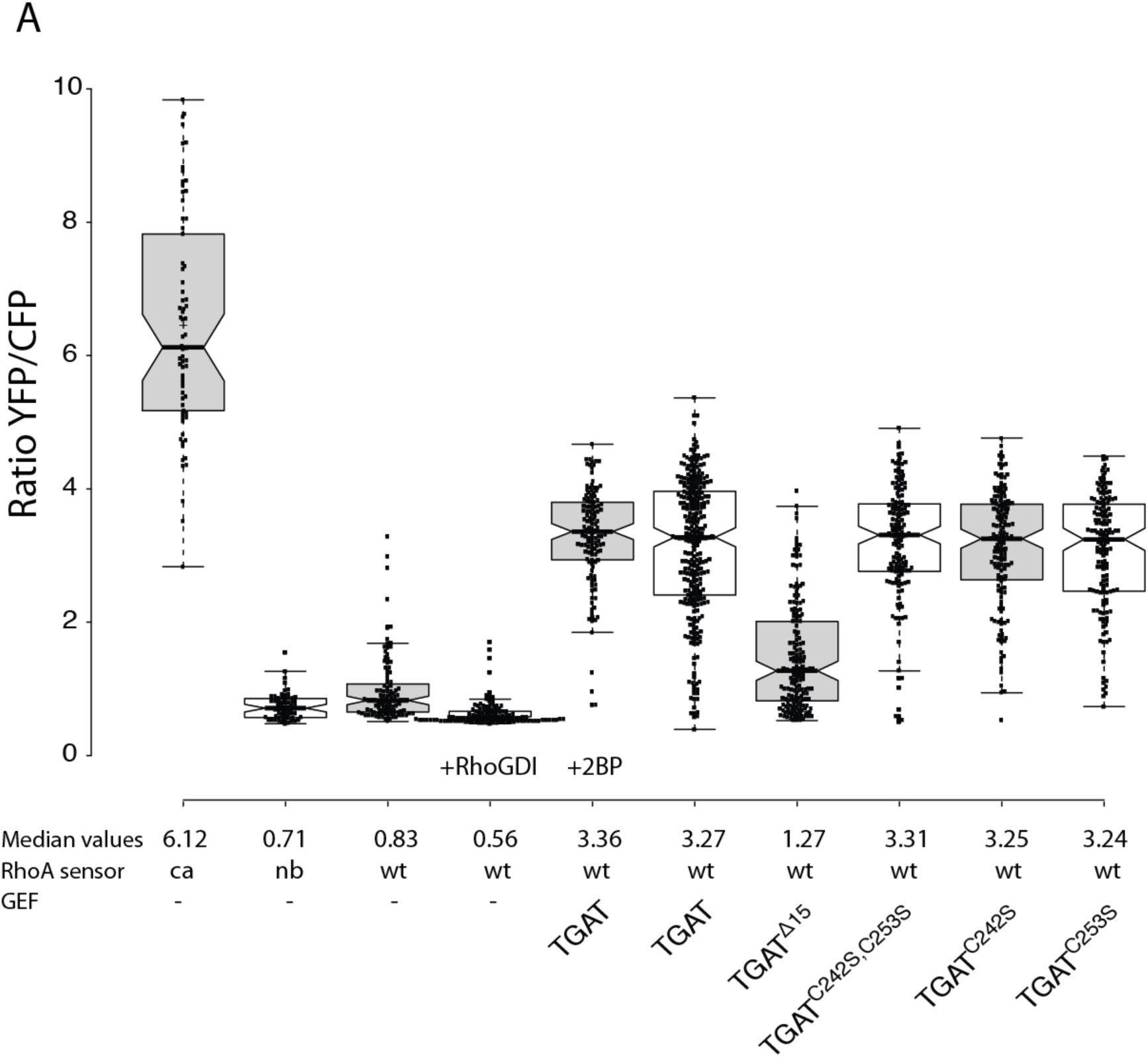
Basal activation of RhoA by the different TGAT mutants. (A) Boxplot showing the median basal YFP/CFP ratio of the DORA RhoA biosensor in HeLa cells. Cells transfected with the constitutive active (ca, *n* = 70) or non-binding (nb, *n* = 62) RhoA biosensor were co-transfected with an empty vector containing only RFP to keep expression levels equal between the different experimental conditions. Wild-type (wt) RhoA biosensor was transfected with an empty vector containing just RFP (control, *n* = 107), RFP-RhoGDI (*n* = 118), RFP-TGAT (+*2-bromopalmitate*) (*n* = 129), RFP-TGAT (*n* = 291), RFP-TGAT^Δ15^ (*n* = 169), RFP-TGAT^C242S, C253S^ (*n* = 148), RFP-TGAT^C242S^ (*n* = 153) or RFP-TGAT^C253S^ (*n* = 150). Boxplot center lines represent the median values; box limits indicate the 25th and 75th percentiles as determined by R software; whiskers extend 1.5 times the interquartile range from the 25th and 75th percentiles; data points from individual cells are plotted as dots.

### The influence of cysteines in the C-tail of TGAT on actin polymerization

Previously, we have shown that plasma membrane located RhoGEF activity results in increased actin polymerization, which was not observed for cytoplasmic located RhoGEF activity (26). In order to investigate the influence of TGAT and its mutants on basal actin polymerization state, HeLa cells were transfected with YFP-TGAT, YFP-TGAT^C242S^, YFP-TGAT^C253S^, YFP-TGAT^C242S, C253S^ or YFP-TGAT^Δ15^. One day after transfection, cells were fixed and stained with an F-actin marker (TRITC-phalloidin) to investigate the influence of the different TGAT mutants on actin polymerization. A clear difference in phalloidin staining intensities was observed between cells transfected with YFP-TGAT and cells transfected with YFP-TGAT^Δ15^ (Figure 5A). Actin polymerization was analyzed quantitatively for the different TGAT variants by normalizing the fluorescence intensities of the phalloidin staining of transfected cells to non-transfected control cells within the same field of view (Figure 5B). These results show that the YFP-TGAT (1.75, 95% CI [1.63 – 1.87]), YFP-TGAT^C242S^ (1.45, 95% CI [1.37 – 1.53]), YFP-TGAT^C253S^ (1.78, 95% CI [1.68 – 1.88]), YFP-TGAT^C242S, C253S^ (1.57, 95% CI [1.50 – 1.64]) conditions all increase actin polymerization compared to TGAT^Δ15^ (1.21, 95% CI [1.12 – 1.30]).

**Figure 5:**
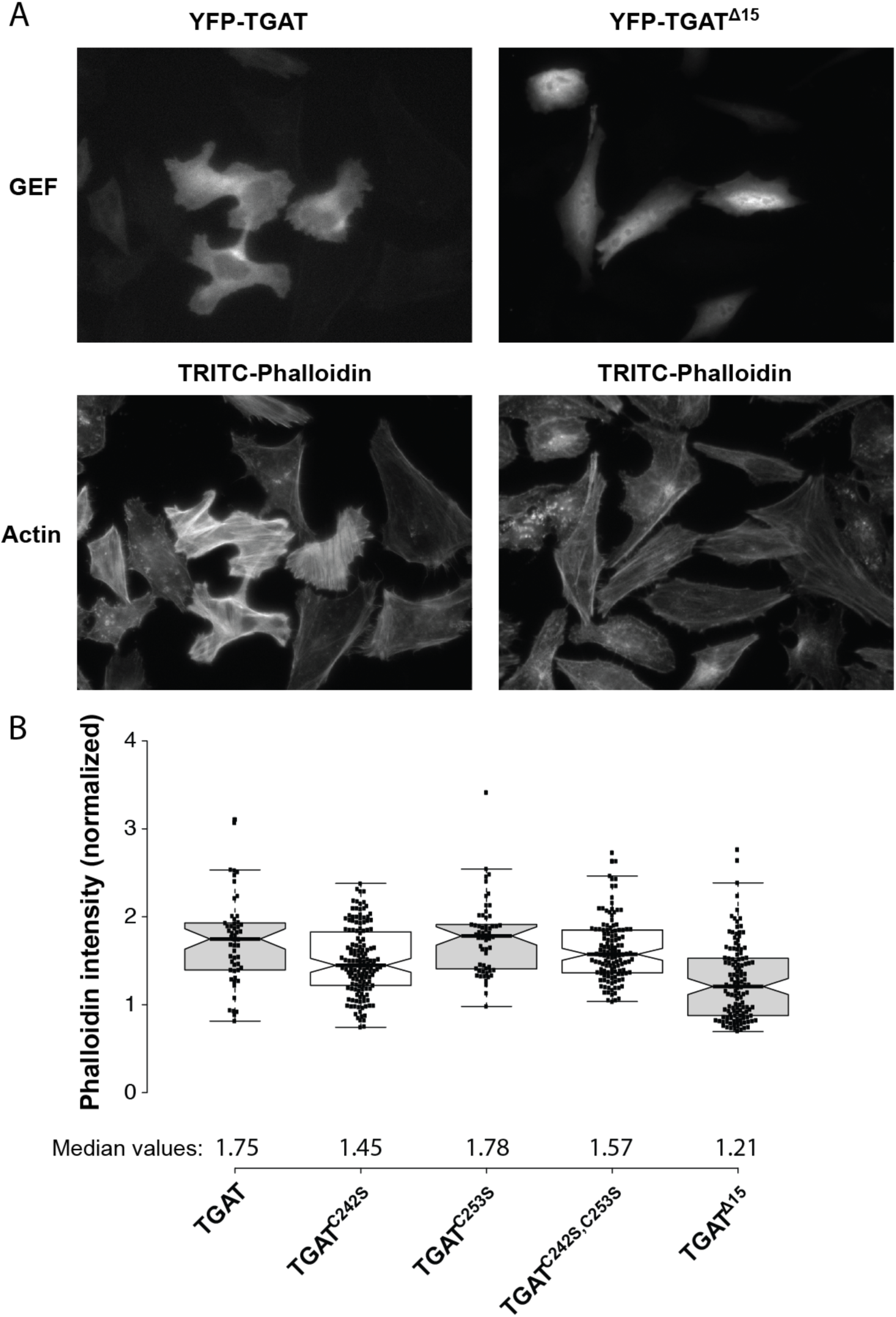
Effect of TGAT mutant expression on actin polymerization. (A) Representative images of HeLa cells transfected with YFP-TGAT or YFP-TGAT^Δ15^ (*top*), stained with TRITC-phalloidin (*bottom*). (B) Quantification of Factin in HeLa cells transfected with YFP-TGAT (*n* = 53), YFP-TGAT^C242S^ (*n* = 147), YFP-TGAT^C253S^ (*n* = 58), YFP-TGAT^C242S, C253S^ (*n* = 128), YFP-TGAT^Δ15^ (*n* = 126) and stained with DAPI and TRITC-phalloidin, as determined by the fluorescent intensity of the TRITC-phalloidin staining. Actin intensity of transfected cells was normalized to the intensity of untransfected control cells in the same field of view.

Another way to quantify actin polymerization in cells is to determine the subcellular localization of the transcription factor Megakaryoblastic Leukemia 2 (MKL2). MKL2 can bind three G-actin molecules through its RPEL motifs, which are released during actin polymerization in cells (29, 30). Upon release of the bound G-actin, MKL2 translocates to the nucleus, making it a bona fide sensor for the G-actin / F-actin status in cells (31). To investigate the influence of the different TGAT variants on the subcellular localization of MKL2, HeLa cells were transfected with MKL2-YFP and the different TGAT variants. Most cells transfected with RFP-TGAT showed a nuclear localization of YFP-MKL2, while YFP-MKL2 was almost always located in the cytoplasm of cells transfected with RFP-TGAT^Δ15^ (Figure 6A). In order to quantify the subcellular localization state of MKL2, the ratio between YFP-MKL2 fluorescence in the nucleus and the cytoplasm was determined for a large number of cells in all conditions (Figure 6B). The median ratio for the RFP-TGAT^Δ15^ condition (1.22, 95% CI [1.12 – 1.32]) was only slightly higher then the median ratio for the control condition (0.96, 95% CI [0.72 – 1.20]) with only a soluble RFP transfected. In contrast, the median ratios for the RFP-TGAT (2.58, 95% CI [2.10 – 3.06]), RFP-TGAT^C242S^ (2.63, 95% CI [2.14 – 3.12]), RFP-TGAT^C253S^ (2.79, 95% CI [2.37 – 3.21]), RFP-TGAT^C242S, C253S^ (3.26, 95% CI [2.73 – 3.79]) were all considerably higher than the control condition (0.96, 95% CI [0.72 – 1.20]) and the RFP-TGAT^Δ15^ condition (1.22, 95% CI [1.12 – 1.32]).

**Figure 6:**
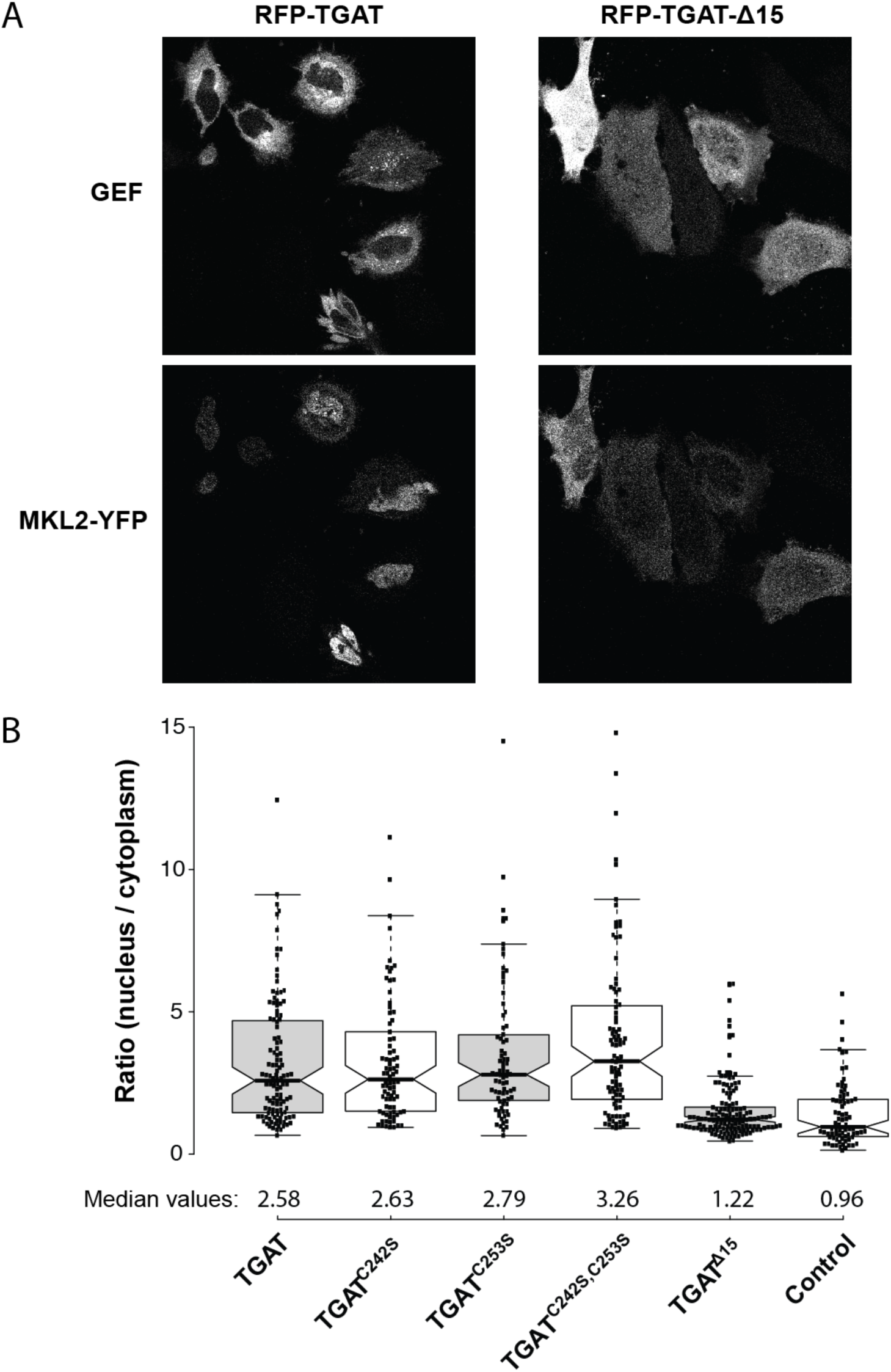
Effect of TGAT mutant expression nuclear MKL2 translocation. (A) Representative images of HeLa cells co-transfected with YFP-MKL2 (*bottom*) and RFP-TGAT or RFP-TGAT^Δ15^ (*top*). (B) Quantification of the ratio between nuclear and cytoplasmic MKL2-YFP fluorescence in HeLa cells transfected with RFP-TGAT (*n* = 116), RFP-TGAT^C242S^ (*n* = 82), RFP-TGAT^C253S^ (*n* = 77), RFP-TGAT^C242S, C253S^ (*n* = 95), RFP-TGAT^Δ15^ (*n* = 142) or RFP (control, *n* = 77).

From the results of the actin staining and the MKL2 localization, we conclude that the cysteine mutants of TGAT do not alter the actin polymerization state of cells compared to wild-type TGAT. In contrast, TGAT^Δ15^ shows a reduced effect on actin polymerization compared to wild-type TGAT. These results fit with the observed differences in RhoA activity, and point towards the critical importance of plasma membrane localization for the function of TGAT.

### TGAT activates RhoA at the plasma membrane and mitochondria

In order to investigate our hypothesis that plasma membrane localization of TGAT results in the activation of RhoA, we decided to use a previously described (26, 32) chemical dimerization system based on rapamycin to gain spatiotemporal control over the subcellular location of TGAT in single living cells. We fused TGAT^Δ15^ to FKBP12 and used FRB fused to several subcellular targeting sequences. RhoA activation was measured over time with the DORA RhoA FRET sensor, before and after targeting FKBP12-TGAT^Δ15^ to Lck-FRB-ECFP(W66A) (plasma membrane), FRB-ECFP(W66A)-CAAX(RhoA) (location of RhoA on endomembranes), MoA-FRB-CFP(W66A) (mitochondria) or Giantin-FRB-CFP(W66A) (golgi apparatus) by adding rapamycin.

Targeting TGAT^Δ15^ to the plasma membrane (Figure 7A) or the CAAX-box of RhoA (Figure 7B) resulted in a fast and sustained increase in RhoA biosensor activation. Interestingly, targeting TGAT^Δ15^ to mitochondria also resulted in a fast and sustained increase in RhoA activation (Figure 7C), whereas targeting TGAT^Δ15^ to the golgi apparatus (Figure 7D) only lead to a minimal response on the RhoA sensor. These results lead us to conclude that, beside the plasma membrane, TGAT has the potential to activate RhoA on several subcellular endomembrane locations as well.

**Figure 7:**
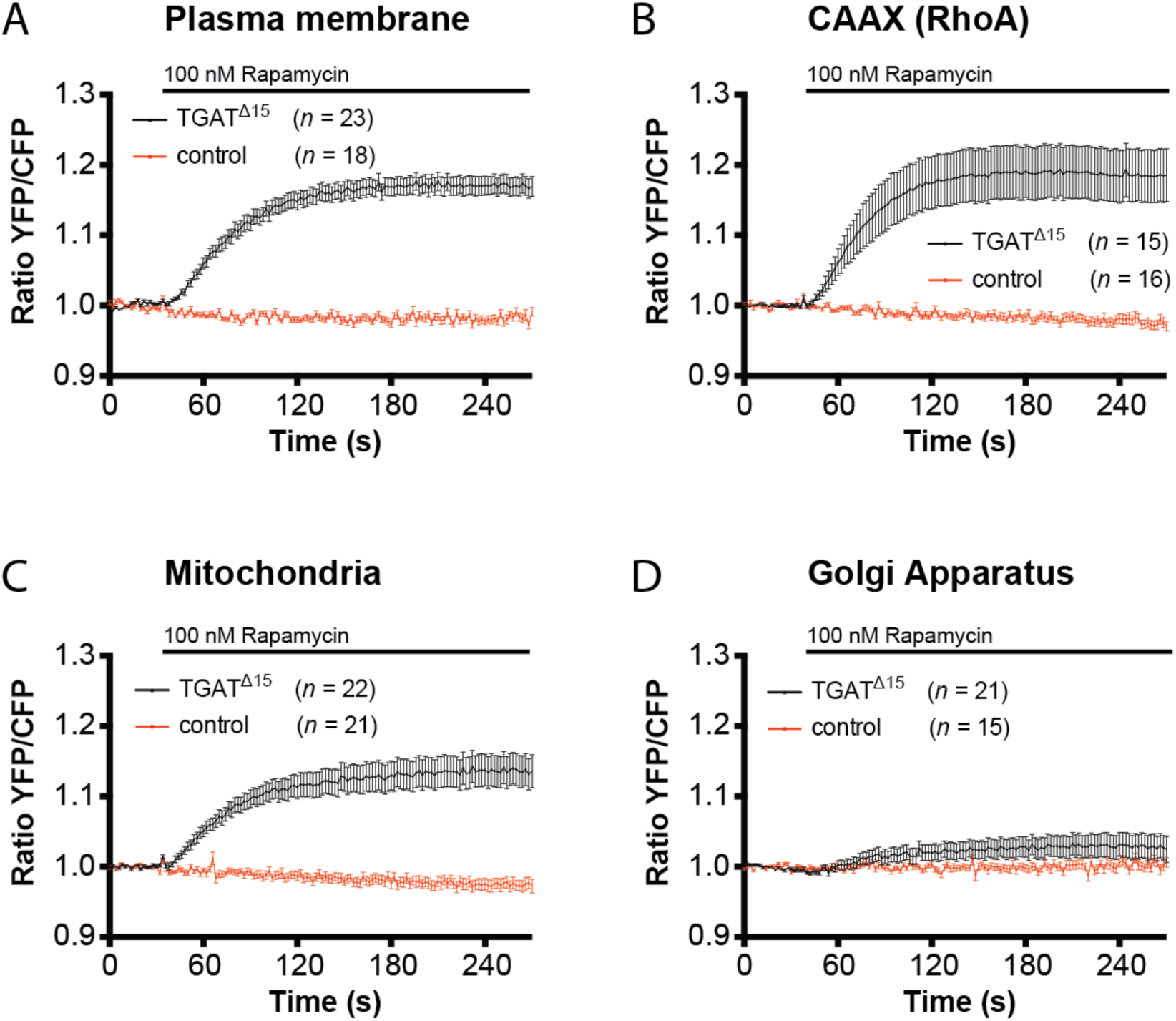
Recruitment of the TGAT^Δ15^ to several subcellular locations results in RhoA activation. (A) Hela cells transfected with the DORA-RhoA biosensor, Lck-FRB-ECFP(W66A) (plasma membrane) and RFP-FKBP12-TGAT^Δ15^ (*n* = 23) or RFP-FKBP12 (control, *n* = 18) were stimulated with Rapamycin (100nM) at *t* = 32s. (B) Hela cells transfected with the DORA-RhoA biosensor, FRB-ECFP(W66A)-CAAX(RhoA) and FKBP12-RFP-TGAT^Δ15^ (*n* = 15) or RFP-FKBP12 (control, *n* = 16) were stimulated with Rapamycin (100nM) at *t* = 32s. (C) Hela cells transfected with the DORA-RhoA biosensor, MoA-FRB-CFP(W66A) (mitochondria) and RFP-FKBP12-TGAT^Δ15^ (*n* = 22) or RFP-FKBP12 (control, *n* = 21) were stimulated with Rapamycin (100nM) at *t* = 32s. (D) Hela cells transfected with the DORA-RhoA biosensor, Giantin-FRB-CFP(W66A) (golgi apparatus) and RFP-FKBP12-TGAT^Δ15^ (*n* = 21) or RFP-FKBP12 (control, *n* = 15) were stimulated with Rapamycin (100nM) at *t* = 32s. Time traces show the average ratio change of YFP/CFP fluorescence (±s.e.m).

## Discussion

Despite the well-established observation that the C-tail of TGAT is essential for its oncogenic potential, it has been unclear what underlying molecular mechanism is responsible. Here, we showed that ectopically expressed TGAT is localized to endomembranes, especially the golgi apparatus. Using a novel quantitative co-localization analysis method for confocal images, we showed that a fraction of wild type TGAT is located at the plasma membrane, while TGAT^Δ15^ is not. Furthermore, we found that mutation of two possible palmitoylation sites in the C-tail, did not affect plasma membrane localization of TGAT. In addition, we observed a reduced mobility of the TGAT mutant devoid of both palmitoylation sites, when compared to mutant TGAT lacking its C-tail, which provides corroborating evidence that this mutant still exhibits membrane affinity. We hypothesized that the three basic residues and 4 hydrophobic residues in the C-tail still provide enough membrane affinity for TGAT to be localized to the plasma membrane. Functional analysis revealed that TGAT and also its cysteine mutants are capable of activating RhoA, resulting in increased actin polymerization.

In contrast, TGAT devoid of the C-tail, TGAT^Δ15^, was severely impaired in the activation of RhoA, actin polymerization or translocation of the transcription factor MKL2. Together these results show that mutation of the cysteines in the C-tail of TGAT does not prevent TGAT from finding its target RhoA on cellular membranes. Synthetic recruitment of TGAT ^Δ15^ from the cytoplasm to various endomembranes demonstrated that TGAT displays enhanced RhoGEF activity when it is present at membranes.

Analogous to our findings, it was previously shown that plasma membrane localization and function of the RhoGTPase Chp is also critically dependent on basic and hydrophobic residues in its C-tail, rather than palmitoylation or prenylation (33). Although the effects of palmitoylation, prenylation and basic or hydrophobic amino acid stretches on endomembrane affinity of proteins have been extensively studied (34–36), it is still unclear how lipidation exactly influences subcellular location and membrane affinity. Whether increase in post-translational lipidation modifications simply provide cumulative gradual increases to endomembrane and plasma membrane affinity, or that specific lipidation motif exist to target proteins to different subcellular endomembrane locations, is still unclear.

One striking outcome of this study is that localization of only a minor fraction of protein at membranes is sufficient to cause substantial changes in cell physiology. Importantly, this membrane localized fraction is almost undetected when employing confocal microscopy under optimal conditions. Only by virtue of a new unmixing-based image analysis strategy, the membrane association can be robustly detected. This method is generally applicable and should be of interest to studies where membrane association is hardly or not visible. Of note, combining the analysis method with high resolution imaging strategies (e.g. SIM or other super resolution techniques) will further lower the limit of detection for membrane association. We found that the mobility of the double mutant was higher than wild-type TGAT, as is expected when proteins spend less time sampling endomembranes due to reduced lipidation. Future site mutagenesis studies targeting the basic and hydrophobic residues in the C-tail could possibly shed more light on whether plasma membrane affinity is specifically involved in its oncogenic potential. Another option would be that the C-tail of TGAT contains unknown motifs for protein-protein interactions or targeting to scaffolds. In any case, our results imply that interfering with palmitoylation is not a viable strategy to reduce oncogenic activity of TGAT towards RhoA. This is in contrast to other GTPases like oncogenic RAS, where it has been postulated that interfering with the depalmitoylation machinery might provide therapeutic benefits by mislocalizing RAS activity (37).

The observation that TGAT can activate RhoA when targeted to mitochondrial sites is unexpected. It was previously shown that the DH domain of p63RhoGEF, which shares 70% sequence identity with the DH domain of TGAT at the protein level, does not activate RhoA at mitochondria in a similar assay. Currently, we do not have an explanation for the increased RhoGEF activity that is observed by mitochondrial localized TGAT-DH.

In summary, our results highlight a role for the C-terminal 15 amino acids in the subcellular location of TGAT and we propose that palmitoylation of the C-tail is dispensable for plasma membrane localization, and therefore activation of RhoA and actin polymerization, of TGAT. Furthermore, we introduce a novel co-localization analysis method for confocal images, which can be used to detect minimal fractions of proteins localized at the plasma membrane. This study provides a framework to further investigate the exact origin of the oncogenic potential in the C-tail from the RhoGEF TGAT.

## Methods

### Construction of fluorescent protein fusions

EGFP-C1-TGAT was a kind gift from Susanne Schmidt (21). mCherry-C1-TGAT was obtained by cutting the EGFP-C1-TGAT vector with XhoI and AgeI and replacing the EGFP for mCherry, cut from mCherry-C1 with the same enzymes. Restriction sites and oligonucleotide overhangs are marked in bold in primer sequences.

To obtain mCherry-C1-TGAT^Δ15^, we performed PCR with EGFP-C1-TGAT as template, by amplifying with forward 5’-GCGCGATCACATGGTCCTG-3’ and reverse 5’-TTT**GGTACC**TCAGGCTACGATTTTCCCGTC-3’. TGAT^Δ15^ was ligated into mCherry-C1 by cutting the vector and PCR product with KpnI and HindIII. To obtain EGFP-C1-TGAT^C242S^, we performed a mutagenesis PCR with EGFP-C1-TGAT as template, by amplifying with forward 5’-GCGCGATCACATGGTCCTG-3’ and reverse 5’-CTCTTTGAACCGATGGCTCAGGGCTACGATTTTCC-3’. mCherry-C1-TGAT^C242S^ was obtained by cutting the EGFP-C1-TGAT^C242S^ vector with HindIII and AgeI and replacing the EGFP for mCherry, cut from mCherry-C1 with the same enzymes.

To obtain mCherry-C1-TGAT^C253S^ and mCherry-C1-TGAT^C242S, C253S^, we performed a PCR with EGFP-C1-TGAT and EGFP-C1-TGAT^C242S^ as template, respectively, by amplifying with forward 5’-AGGTCTATATAAGCAGAGC-3’ and reverse 5’-TT**GGTACC**TCAAAACCAACTAATTTCACGAAAAGTCTCTTTG-3’. TGAT^C253S^ and TGAT^C242S, C253S^ were ligated into mCherry-C1 by cutting the vector and PCR products with Acc651 and HindIII.

CFP and YFP color variants were obtained by cutting mCherry-C1-TGAT, mCherry-C1-TGAT^Δ15^, mCherry-C1-TGAT^C242S^, mCherry-C1-TGAT^C253S^ and mCherry-C1-TGAT^C242S, C253S^ with AgeI and HindIII and replacing the mCherry with mTurquoise1 or mVenus, cut from mTurquoise1-C1 and mVenus-C1 with the same enzymes.

For the rapamycin experiments, mCherry-FKBP12-C1-TGAT^Δ15^ was created by cutting mCherry-C1-TGAT^Δ15^ with AgeI and HindIII and replacing mCherry with FKBP12-mCherry, cut from FKBP12-mCherry-C1 (26) with the same enzymes. A membrane targeting sequence (derived from amino acid residue 1-10 of Lck; MGCVCSSNPE) was constructed by annealing (38) two oligonucleotide linkers, 5’-ctagccaccatgggctgcgtgtgcagcagcaaccccgagcta-3’ and 5’-ccggtagctcggggttgctgctgcacacgcagcccatggtgg-3’, with sticky overhangs and inserting it into an mVenus-C1 plasmid cut with NheI and AgeI, resulting in Lck-mVenus. Lck-mTurquoise2 was obtained by exchanging mVenus for mTurquoise2 in the Lck-mVenus plasmid by cutting with AgeI and BsrGI. The Lck-FRB-ECFP(W66A) was a kind gift from M. Putyrski (39). FRB-ECFP(W66A)-Giantin, ECFP(W66A)-FRB-MoA, mVenus-MKL2 and the DORA RhoA sensors were previously described (26). In order to obtain mTurquoise2-CAAX(RhoA), two oligonucleotides were annealed as previously described (38). Annealing forward 5’-

**GTAC**aagctgcaagctagacgtgggaagaaaaaatctgggtgccttgtcttgtga**G**-3’ and reverse 5’-**GATC** ctcacaagacaaggcacccagattttttcttcccacgtctagcttgcag**CTT**-3’ oligonucleotides yielded the coding sequence for the last 15 amino acids of the C-terminus of RhoA (LQARRGKKKSGCLVL*) with overhangs (in capitals) on both sides, compatible with BsrGI and BamHI restriction sites. The double stranded linker was ligated into a C1-mTurquoise2 vector cut with BsrGI and BamHI, resulting in mTurquoise2-CAAX(RhoA). FRB-ECFP-CAAX(RhoA) was obtained by ligating the CAAX(RhoA) fragment, cut from mTurquoise2-CAAX(RhoA), into FRB-ECFP(W66A)-Giantin cut with the same enzymes. mRFP-RhoGDI was a kind gift from Martin A. Schwartz (40). Plasmids constructed in this study will be made available through Addgene: http://www.addgene.org/Dorus_Gadella/.

### Cell Culture & Sample Preparation

HeLa cells (American Tissue Culture Collection: Manassas, VA, USA) were cultured using Dulbecco’s Modified Eagle Medium (DMEM) supplied with Glutamax, 10% FBS, Penicillin (100 U/ml) and Streptomycin (100µg/ml). Cell culture, transfection and live cell microscopy conditions were previously described (26).

### Widefield microscopy

Static and dynamic ratiometric FRET measurements in HeLa cells were performed using a wide-field fluorescence microscope (Axiovert 200 M; Carl Zeiss GmbH) kept at 37°C, equipped with an oil-immersion objective (Plan-Neofluor 40×/1.30; Carl Zeiss GmbH) and a xenon arc lamp with monochromator (Cairn Research, Faversham, Kent, UK). Images were recorded with a cooled charged-coupled device camera (Coolsnap HQ, Roper Scientific, Tucson, AZ, USA). Typical exposure times ranged from 50ms to 200ms, and camera binning was set to 4x4. Fluorophores were excited with 420 nm light (slit width 30nm) and reflected onto the sample by a 455DCLP dichroic mirror, CFP emission was detected with a BP470/30 filter, and YFP emission was detected with a BP535/30 filter by rotating the filter wheel. All acquisitions were corrected for background signal and bleedthrough of CFP emission in the YFP channel (around 55% of the intensity measured in the CFP channel). In dynamic experiments, cells were stimulated with 100nM Rapamycin at the indicated time points (LC Laboratories, Woburn, USA).

In the actin staining experiment, DAPI was excited with 420 nm light (slit width 30nm) and reflected onto the sample by a 455DCLP dichroic mirror and emission was detected with a BP470/30 filter, YFP was excited with 500nm light (slit width 30nm) and reflected onto the sample by a 515DCXR dichroic mirror and emission was detected with a BP535/30 filter. RFP was excited with 570 nm light (slit width 10nm) and reflected onto the sample by a 585CXR dichroic mirror and emission of RFP was detected with a BP620/60 filter.

### Confocal microscopy

Experiments were performed using a Nikon A1 confocal microscope equipped with a 60x oil immersion objective (Plan Apochromat VC, NA 1.4). For the co-localization experiments the pinhole size was set to 1 Airy unit (<0.8µm) and images with 1.5x zoom of 1024x1024 pixels were acquired. For the MKL2 translocation experiments, the pinhole size was set to 1 Airy unit (<0.8µm) and images were acquired with 1x zoom using tile scans, resulting in images of 4660x4660 pixels. Samples were excited with 447nm, 514nm and a 561nm laser line, and reflected onto the sample by a 457/514/561 dichroic mirror. CFP emission was filtered through a BP482/35 emission filter; YFP emission was filtered through a BP540/30 emission filter; RFP emission was filtered through a BP595/50 emission filter. All acquisitions were corrected background signal. To avoid bleed-through, images were acquired with sequential line scanning modus.

### Fluorescence correlation spectroscopy

Cells were transfected with 100ng of mTurquoise1-TGAT, mTurquoise1-TGAT^C242S, C253S^ and mTurquoise1-TGAT^Δ15^ or mTurquoise1 as control. FCS data was acquired at an Olympus FV1000 confocal microscope equipped with a Picoharp TCSPC module (Picoquant, Germany). Sample were mounted on the table and illuminated with a pulsed 440nm Picoquant diode laser (20 MHz, 0.6 kW.cm^-2^) using an Olympus UPLS Apo 60x water NA1.2 objective lens. The fluorescence signal was detected for 60-120s in confocal mode with the pinhole diameter set at 130μm. The fluorescence passed a 440 dichroic mirror, was filtered by a 460-500 nm emission filter and detected by an avalanche photodiode (MPD). Correlation curves were generated in FFS Dataprocessor (v2.3 SSTC, Belarus) and fitted using a triplet state-diffusion model (41). Since the TGAT fusion proteins can be present as free cytoplasmic protein and bound to the membrane, two diffusion times were included in the fitting model. The diffusion times were globally linked over the various measurements, the volume shape factor was fixed to the value obtained from the calibration sample and the triplet time was restricted between 0 and 50μs. To compare the diffusion times between the various measurement days the values were converted into diffusion coefficients (D) (41), taking into account the slightly variable sizes of the detection volume from day to day, calibrated by the mTq1 sample (*D* = 90 μm^2^.s^-1^ in PBS). The average diffusion coefficient was calculated taking into account the fractions and values of the two retrieved diffusion coefficients.

### Actin staining

HeLa cells transfected with different TGAT constructs were washed with phosphate-buffered saline solution (PBS) and fixed with 4% formaldehyde for 20 minutes. After washing with PBS, cells were permeabilized with PBS containing 0.2% Triton X-100. After a second wash step with PBS and blocking of non-specific binding by 1% BSA in PBS for 10 minutes, cells were stained with 0.1µM TRITC-phalloidin (Sigma-Aldrich) and 0.1µg/ml DAPI. After washing with PBS, cells were mounted in Mowiol and fluorescence images were attained using a widefield fluorescence microscope (Axiovert 200 M; Carl Zeiss GmbH).

### Image Analysis

ImageJ (National Institute of Health) was used to analyze the raw microscopy images. Further processing of the data was done in Excel (Microsoft Office) and graphs and statistics were conducted using Graphpad version 6.0 for Mac, GraphPad Software, La Jolla California USA, <u>www.graphpad.com</u>. Boxplots in Figure 3, Figure 4, Figure 5 and Figure 6 were generated online, using the website <u>http://boxplot.tyerslab.com/</u>. For the MKL2 transcription factor experiments, MKL2 intensity in cytoplasm and nucleus were measured and their ratio was determined. The static FRET data in Figure 4 was processed using a custom made MatLab GUI, which was described before (26).

The confocal image analysis of the plasma membrane localization for the different constructs and controls was performed by using a combination of ImageJ and MatLab scripts (MATLAB, The MathWorks, Inc., Natick, Massachusetts, United States). For each construct confocal images were obtained with a CFP, YFP and RFP channel (1024×1024 pixels with pixel size 140 nm). A sequential time series of 8 images was recorded, which were subsequently averaged for each channel. The channels were spatially registered based on a shift determined with the Lucas-Kanade method that was performed on the controls. The positive control, i.e. Lck-CFP, Lck-YFP and soluble RFP was used to determine the spatial shift between the CFP and the YFP channel. The negative control, i.e. Lck-CFP, soluble YFP and RFP was used to determine the spatial shift between the RFP and YFP channel. In each image and channel the background was determined and subtracted from the image, all sets from one construct were subsequently stored as an ImageJ tif ‘hyperstack’. In ImageJ lines (10 px wide and ~6-10 µm long) were drawn perpendicular on regions with a well-defined cytoplasm-plasma membrane-extracellular transition, whilst carefully switching between channels to avoid inclusion of spurious structures in the line-scan. Lines were drawn in all possible orientations, in order to avoid any possible bias resulting from imperfect registration. For each image the line-scans were stored as a RoiSet. By using a matlab script the RoiSets from ImageJ were imported and line-scans were performed in Matlab using bilinear interpolation. In one set, all the profiles obtained from the line-scans for all channels were aligned and centered on the plasma membrane peak position in the Lck-CFP channel, and the profiles were oriented in the same manner, i.e. cytoplasm on the left hand side. By using bilinear interpolation all the profiles were placed on the same position axis. The average profiles were calculated for all three channels and any residual background observed in the extracellular space (>2 µm) was subtracted, after which the channels were normalized to unity based on the cytoplasm region ([−2.5, −1.5] µm). Because the Lck-CFP channel also contained a cytoplasmic localization, the cytoplasmic component was subtracted in order to obtain a more accurate plasma membrane component. This corrected Lck-CFP (*f*_*(CFP-RFP)*_) and the RFP (*f*_*RFP*_) profiles were subsequently used to linearly unmix the profile from the YFP channel (*f*_*YFP*_), using constrained linear regression (lqlin) within the region [-2.25, 2.25] µm. Hence, *f*_*YFP*_ = *a*_*CP*_ *f*_*RFP*_+*a*_*PM*_ *f*_*(CFP-RFP)*_, where the coefficients *a*_*CP*_ and *a*_*PM*_ were constrained to positive values and correspond to the cytoplasmic and plasma membrane component respectively. All profiles were realigned and centered on the unmixed PM component. The residual peak in the negative control condition (3 ± 1%) is likely to be caused by the difference in the points spread function (PSF) between YFP and RFP. The PSF of RFP is slightly wider than that of YFP, which has a smoothing effect on the mCherry profile. This is also consistent with the shape of the profiles in Figure 6B and the residuals (not shown). In order to estimate the confidence intervals on the unmixed CP and PM profiles a bootstrap was performed. Random sets (*n* = 1000) were drawn from the original set with replacement, and the same normalization and unmixing was performed on these sets, after which 95% confidence interval was calculated based on the standard error of the mean.

## Acknowledgments

We thank Roy Baas for his contribution to the construction of the Lck-mVenus plasmid.

## Author contributions

J.v.U, J.G, M.H, and D.B. performed experiments, analyzed the data and wrote the manuscript. Y.I.W. and T.Y designed and constructed the DORA-Rho GTPase biosensor. M.P developed and performed image analysis procedures. T.W.J.G. assisted with experimental design and interpretation of data. All authors approved the final manuscript.

